# Morphofunctional analysis of antigen uptake mechanisms following sublingual immunotherapy with beads in mice

**DOI:** 10.1101/370163

**Authors:** Yaser Hosny Ali Elewa, Tatsuya Mizoguchi, Osamu Ichii, Teppei Nakamura, Yasuhiro Kon

## Abstract

**Background:** Recently, sublingual immunotherapy (SLIT) has been used as a safe and efficient method for the treatment of and immunization against asthma and various allergies. However, the routes of antigen uptake through the mucosa of the oral cavity remain incompletely understood, as do the roles of sex and age in the process. For this purpose, to elucidate the mechanism and efficacy of SLIT among different sexes and ages microbeads were dripped into the sublingual region to mimic antigen uptake by the sublingual mucosa.

**Methods:** Twenty microliters of either phosphate buffered saline (PBS) or fluorescently labelled microbeads (latex and silica beads) were placed under the tongue of both male and female C57BL/6 mice at young (3 months) and old (6 months) ages. The lower jaw was examined 30 min after administration, and beads were detected with a fluorescence stereomicroscope. Morphological observations of the mucosa of the fluorescent areas were made with scanning electron microscopy (SEM) and an all-in-one light fluorescence microscope (LM). Fluorescence intensity was compared between both sexes and ages.

**Results:** Stereomicroscopic observation revealed fluorescent illuminations in three compartments of the sublingual mucosa: the sublingual caruncles (SC), the oral rostral mucosa (OR) and the buccal mucosa (BM). Interestingly, the fluorescence intensity tended to be higher among females than among males in the SC region in particular. However, there were no significant age-related differences. SEM and LM revealed beads in the lumina of both mandibular ducts and sublingual ducts (Sd). Additionally, the apical cytoplasm of some Sd cells contained silica beads. However, there were no specification in the OR mucosa or BM.

**Conclusions:** This study reveals the major role Sd play in local immunity via the antigen uptake mechanisms. Furthermore, our data suggest that the efficacy of SLIT in humans could be affected by sex.

## INTRODUCTION

In both humans and animals, the prevalence of allergic diseases such as seasonal rhinitis and atopic dermatitis has increased substantially in recent decades [1-3]. The symptoms accompanying such allergic conditions range in severity. Mild symptoms such as itching and sneezing may cause disturbances in the patient’s daily life and affect productivity, while severe ones such as anaphylactic shock can be life-threatening [4]. Therefore, establishing countermeasures against the development of allergic conditions is an important issue in both the medical and veterinary fields.

Treatment for allergic diseases is currently based primarily on symptomatic therapy to reduce inflammation; antihistamines and steroids are widely used for this [5, 6]. However, immune induction therapy has attracted attention in recent years. Among them is sublingual immunotherapy (SLIT), which is allergen-specific. In SLIT, the allergens or antigens are administered to the lower part of the tongue and may provide sustained and safe therapeutic effects [7-9]. With this method, the amount of antigen administered to the lower part of the tongue is gradually increased to induce immune tolerance and improve the patient’s hypersensitivity symptoms [10]. Interestingly, because SLIT administration and postoperative management are so straightforward, recently there has been increasing interest in the clinical application of SLIT in humans [11], as well as in the treatment of atopy and mite allergies in dogs [12].

Antigen uptake through the sublingual mucosa following SLIT is the mechanism by which antigen-specific immune tolerance is induced. Therefore, immunological and morphologic functional evaluations of the oral mucosa are essential for further characterizing this mechanism. From an immunological point of view, several previous reports have suggested that antigen-presenting cells “APCs” (dendritic cells, macrophages) and regulatory T-cells in the sublingual mucosa play a role in antigen uptake and induction of immune tolerance following SLIT [13-17]. Furthermore, a recent report revealed the role of APCs in sublingual ductal epithelial cells in the transportation of sublingual antigen. This was shown using soluble antigens such as ovalbumin and particulate antigens such as E. coli, latex beads (Lt) and silica beads (Si) [18]. Moreover, it has been revealed that bacterial infection of the salivary glands may result from bacteria ascending through salivary gland ducts and stasis of salivary flow through the ducts [19]. This suggests that salivary gland ducts play a role in antigen uptake. After morphological analysis of the different compartments of the oral mucosa, their roles in antigen uptake remains unclear. Interestingly, sexual dimorphism of the rodent submandibular gland granular duct (granular convoluted tubule) has been reported [20]. Further, it has been revealed that aging affects the structure of salivary glands [21]. However, there have been no reports regarding differences in the therapeutic efficacy of SLIT between patients of different sexes or ages. Therefore, in this study we morphologically analyzed the bead accumulation sites in the oral cavity mucosa of young and old mice of both sexes following sublingual administration of either Lt or Si. The accumulation sites reflect the specific anatomic regions of antigen uptake. We found that the bead-derived fluorescence was mainly observed in the dorsal part of the sublingual caruncle (SC), as well as in the oral rostral (OR) and buccal mucosa (BM) of the oral cavity proper.

Interestingly, the fluorescence intensity was higher in the SC and OR mucosa than that in the BM. Further, the females tended to demonstrate higher intensity than males. In the Lt but not Si beads group, significant sex differences were observed, especially in the SC region. However, no significant age-related changes were observed in either bead group. We suggest that the localization of the beads could be affected by the sex and the material of the bead. Furthermore, we suggest that the dorsal part of the SC may play an important role in local immunity related to antigen uptake mechanism.

## MATERIALS AND METHODS

### Ethics Statement

The investigators conducted experiments in accordance with the guidelines for the Care and Use of Laboratory Animals, Hokkaido University, Graduate School of Veterinary Medicine (certified by the Association for Assessment and Accreditation of Laboratory Animal Care International). All experiments were conducted according to the protocols approved by the Institutional Animal Care and Use Committee of the Graduate School of Veterinary Medicine, Hokkaido University, Japan (approval No. 15-0079).

### Experimental animals and experimental design

Male and female C57BL/6N (B6) mice of both young age (12-16 ws) and old age (24-36 ws) were purchased from Japan SLC (Hamamatsu, Japan) and were used for each experiment. They were anesthetized via intraperitoneal administration of a mixture of medetomidine (0.3 mg/kg), midazolam (4.0 mg/kg) and butorphanol (5.0 mg/kg). Subsequently, male and female mice of each age received 20 μL of either phosphate buffered saline (PBS) (control groups) or 1% fluorescence-labelled beads suspended in PBS (experimental groups) on the lower part of the tongue after the tongues were raised with tweezers (S1a Fig.). The experimental groups were subdivided into a latex (Lt) group, which received latex beads (Fluoresbrite ™ YG carboxylate microsphers, diameter 0.75 μm, 1% Polyscience, Warrington, PA., USA), and a silica (Si) group, which received silica beads (diameter 0.8 μm, 50 mg/mL; Micromod Partikel technologie GmbH, Warnemurende, Germany). Thirty min after the beads were applied, the common carotid artery was cut, and the mice were euthanized by exsanguination. The lower jaw was then separated from the upper one and washed with 0.01 M PBS.

### For stereoscopic microscopic observation

The sublingual mucosa was examined with a stereomicroscope after cutting the free tip of the tongue. The sites of fluorescently labelled bead accumulation were observed and photographed using the fluorescent stereomicroscope (AXIO ZOOM-V 16, ZEISS, Tokyo, Japan) for all experimental groups and compared with the control group.

### Tissue preparation for light microscopic observation

Following fixation of the lower jaw in either 10% neutral buffered formalin at 4 °C for 24 hr or 4% paraformaldehyde (4 °C, overnight), the samples were decalcified in formic acid at room temperature for 2 days. After washing the samples in PBS three times, 5 min each, the sublingual region was separated just caudal to the SC region under the stereoscopic microscope. The sublingual region was then dehydrated in ascending grades of alcohol, and embedded in paraffin. Three μm serial paraffin sections of the sublingual region were prepared, stained with hematoxylin and eosin (HE). The stained sections were then observed and photographed using BZ-X710 all-in-one fluorescence microscope (Keyence, Osaka, Japan).

### Tissue preparation for scanning electron microscopy

In the silica bead and control groups, the lower jaw was fixed with 2.5% glutaraldehyde (4 °C, overnight). Thereafter, it was immersed six times in a 0.1 M PBS (pH 7.4) for 10 minutes each. Subsequently, at 4 °C, the samples were rinsed in a 0.5% tannic acid solution for 10 minutes and 1.0% tannic acid solution for 1 hour, and then in 0.1 M phosphoric acid. After washing in PBS for 15 min, the specimens were dehydrated stepwise using a series of graded ethanol. After dehydration, the samples were transferred into a mixture of 100% ethyl alcohol and isoamyl acetate (1:1), then kept for 20 min twice in isoamyl acetate solution, and dried in a critical point drier (HCP-2, Hitachi, Tokyo, Japan). Thereafter, the dried samples were fixed on aluminum stubs with double-faced adhesive tabs. Surface treatment with a 20-nm thick platinum layer was carried out with ion sputtering (E-1030, Hitachi, Tokyo, Japan) for 1 min. The samples were then observed and photographed using a scanning electron microscope (SU 8000 field emission scanning electron microscope, Hitachi, Tokyo, Japan, conditions of 10 kV).

### Histoplanimetry

To calculate the fluorescence detection rate within different sublingual compartments of both the Lt and Si bead groups, the number of samples in which fluorescence was observed in the SC, OR, and BM mucosa was calculated and divided by the total number of analyzed samples. Moreover, for evaluation of the fluorescence intensity as an indicator of bead accumulation, images from the experimental group samples were captured. Following this, the photographed image was monochromatized using JTrim (free software, manufactured by Woody Bells, Japan). Thereafter, the luminance was measured ten times (ImageJ; NZH, Bethesda, MD, USA) at each site of the SC, OR and BM. The average value was quantified and defined as the fluorescence intensity. This value was then compared with the control non-fluorescent area.

### Statistical analysis

Significant difference of the fluorescence intensities in the measurement sites (SC, OR, BM) when compared to that of negative control area in each group was analyzed by the Dunnett t method after the Kruskal-Wallis test. The differences of the fluorescence intensities between groups were compared using the Scheffé’s method after the Kruskal–Wallis test. The analysis of the gender and age group differences was applied by Mann-Whitney *U* test. In all analysis, a *P* value < 0.05 was regarded as a significant difference.

## RESULTS

### Morphological observation of sublingual mucosa in mouse lower jaw following PBS or bead administration

In the experimental group, the fluorescently labelled bead accumulations in the sublingual mucosa were mainly observed in three sites: the caudal protrusion that represent the SC; two lateral sides that represent the BM; and at a depression in the median rostral position just behind the incisor teeth that represents the OR and appeared as an elliptical fluorescence accumulation site. An area with weak fluorescent illuminations cranial to the SC was used as a negative control area (S1b and c Figs.). With SEM, the SC appeared as a paired mucosal protrusion of average width 400 μm. The BM appeared as two lateral grooves extending toward the rostral side at the boundary between the buccal mucosa and the bottom of the oral cavity proper. The OR appeared as a median depression (about 150 μm × 700 μm) on the caudal side of the lower incisor teeth (S1c Fig.). In the control group, no fluoresce could be observed by fluorescent stereomicroscope (S1d Fig.).

### Fluorescence detection rates and intensities in SC, BM, and OR among young and old ages of both sexes

As shown in Table 1, we compared the fluorescence detection rates in the SC, BM, and OR sites in young and old groups of males and females in both Lt and Si bead experimental groups. In the Lt bead experiment group, the fluorescence was detected at a rate as high as 100% of the average detection rate of all groups in the OR site. The average detection rate in the SC was 87.5%. The BM had lower values than the other two sites (69%). The old female group had a detection rate of 100% at all three sites. In the Si bead experimental groups, the young and old female group showed 100% detection rate in the OR and the SC. The detection rate in the old males group was 75%. Furthermore, the average fluorescence detection rate in the BM among all groups was lower than the other two sites (50%).

**Table 1.** Fluorescence detection rate (percentage) of the both Si and Lt beads among different sites, ages, and sexes.

|  | Sublingual |  | Buccal |
| --- | --- | --- | --- |
| **A- Latex bead group** | caruncle | Oral rostral | mucosa |
| Young male | 75.0 | 100.0 | 50.0 |
| Young female | 100.0 | 100.0 | 75.0 |
| Old male | 75.0 | 100.0 | 50.0 |
| Old female | 100.0 | 100.0 | 100.0 |
| **Average** | 87.5 | 100.0 | 68.8 |
| **B- Silica bead group** |  |  |  |
| Young male | 100.0 | 100.0 | 50.0 |
| Young female | 100.0 | 100.0 | 50.0 |
| Old male | 75.0 | 75.0 | 25.0 |
| Old female | 100.0 | 100.0 | 75.0 |
| **Average** | 93.8 | 93.8 | 50.0 |

Additionally, we analyzed the fluorescence detection and intensities in both Lt and Si bead groups among young and old groups, as well as among male and female (Fig. 1). In Lt bead administration, the females in both young and old groups tended to exhibit stronger fluorescence at all sites (SC, OR, and BM) than males did (Figs. 1a-d). Interestingly, the SC in females showed stronger fluorescently labelled bead accumulations (Figs. 1b, and d) than those in males (Figs. 1a, and c) in both age groups.

**Fig. 1.**
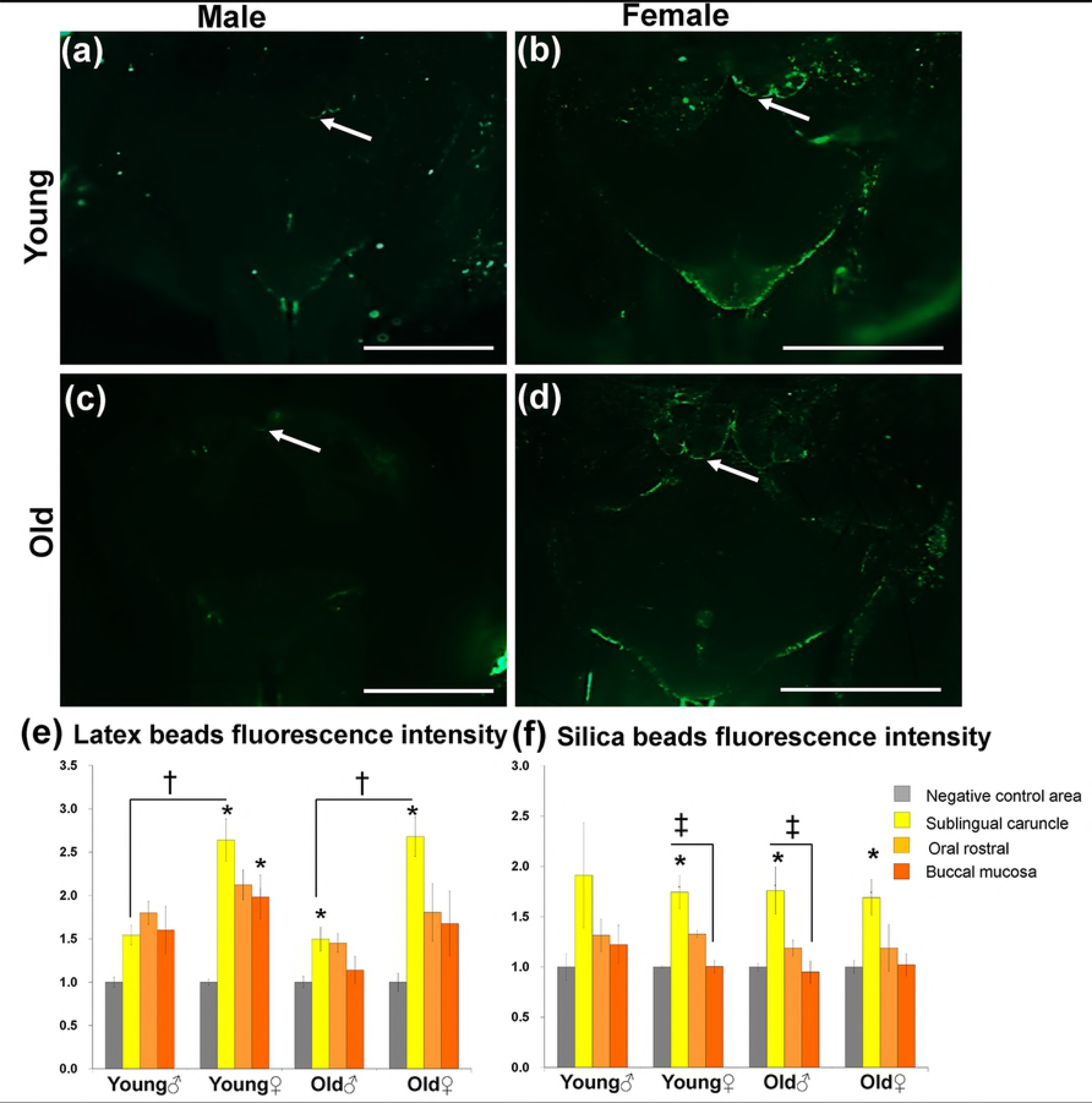
Localization of fluorescent beads in the oral mucosa of the lower jaw in the latex beads group (a-d). Fluorescence microscopy observation in both the young group (a and b) and old group (c and d). Notice stronger fluorescence in the young and old female groups than in male groups in particular for SC (white arrow). Scale bars = 1 mm. Graphs showing the fluorescence intensities in both Lt (e) and Si (f) bead groups. The fluorescence intensity was measured for the SC, OR, BM, and the numerical value was obtained by scoring the negative control value as 1.0. Values are given as the mean ± SE, n= 4. *: Significant difference in the measurement sites (SC, OR, BM) from the negative control in each group (Kruskal-Wallis test, Dunnett t method, P <0.05). †: Significant difference between sexes (Mann-Whitney *U* test), ‡: Significant difference between sites (Kruskal-Wallis test, Scheffé’s method).

However, neither sex nor age-related changes were observed in the OR and BM. On the other hand, weaker fluorescently labelled bead accumulations were observed in the Si bead group than that in the Lt group at all sites. Further, in the Si bead group, the fluorescently labelled bead accumulations tended to be slightly stronger in the SC than other sites (data not shown).

To determine variations in bead accumulations among both the Lt and Si bead groups and among both ages and sexes, the fluorescence intensity was quantified by an image analysis method in all accumulation sites (Figs. 1e, and f) in comparison with the negative control area (reference value = 1.0) (Fig. S1b). In the Lt bead group, the fluorescence intensity in the SC, OR, and BM tended to have a higher value than in the negative control area in both age and sex groups. The intensity had significantly higher values in the SC of the young male group and young female group and the BM of the young female group. Furthermore, significant differences in SC fluorescence intensity were found between males and females in both the young and adult groups (Fig. 1e).

In the Si bead group, as with the Lt group, the SC tended to have a higher value than the negative control area in all groups. Moreover, except for the young male group, the fluorescence intensity in the SC was significantly higher than in the negative control area. Additionally, the fluorescence intensity in the SC of the young female and old male groups was significantly higher than in the BM. On the other hand, there was no sex difference as in the Lt bead group (Fig. 1e). No significant age-related changes were observed in both bead groups (Figs. 1e, and f).

### SEM and light microscopic observations of the structure of sublingual mucosa in mouse lower jaw

SEM observation of the mucosal surface morphology among both sexes revealed that there were no remarkable differences between sexes. The SEM observation revealed that the SC in the control group (female, 3 months old) appeared as a pair of right and left protrusions. A groove was observed in the central part of the protrusion extending toward the rostral side. In addition, a deep depression (approximately 30 μm long and 10 μm short) was seen on the dorsal surface of the caudal side of the SC (Figs. 2a, b). The dorsal surface of this depression was smooth; however, the surface of the edge that bordered the depression showed fine folds (Fig. 2c). The SEM of the dorsal surface of the SC in the Si bead group revealed numerous beads around the depression on the dorsal surface of the caudal side of the SC and along the groove extending in the SC (Figs. 2d-f). The Si beads were observed as spherical particles of well-defined size (Fig. 2g).

**Fig. 2.**
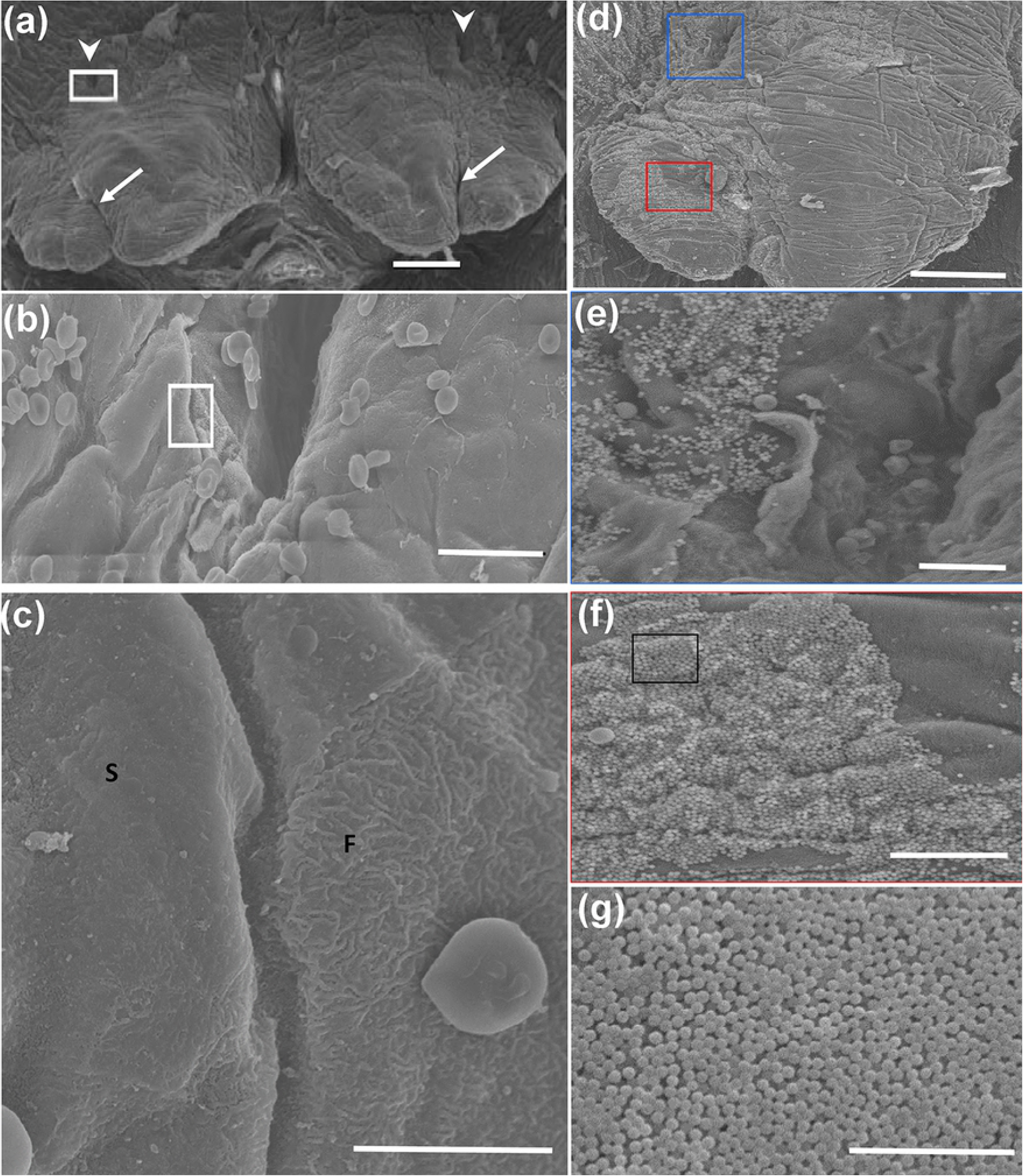
SEM of the dorsal surface of the SC of the control group (female, 3 months old) (a-c) and silica bead group (d-g). (a) The SC are observed as a pair of left and right protrusions. Notice the central grooves (arrows) and recessed dorsal aspect (arrow heads). Scale bar = 100 μm. (b) Higher magnification of the white framed area in (a). The entrance to the recess can be seen in the center. Notice the boundary between the mucosal epithelial cells of the SC and the epithelial cells constituting the edge of the recess (white frame). Scale bar = 20 μm. (c) Higher magnification of the white boxed area in (b). Notice the smooth surface (S) of the mucosal lining the SC and fine folded surface (F) lining the entrance to the recess. Scale bar = 10 μm. (d) SEM of the dorsal surface of the SC of the experimental group (female, 3 months old). Notice the accumulation of silica beads in the periphery around the tip of the SC (red box) and in the recessed part (blue box) on the dorsal side. Scale bar = 100 μm.(e) Higher magnification of the blue boxed area in (d). Notice numerous silica beads adherent to the mucosal surface around the recess. (f) Higher magnification of the red boxed area in (d). (g) Higher magnification of the boxed area in (f). Notice the silica beads with a spherical uniform shape. (d-g) Scale bars = 15 μm.

We then examined the distribution of fluorescently labelled Si beads in H&E stained paraffin sections. The two sublingual and mandibular ducts opened separately into the sublingual mucosa (Figs. 3, and 4a-f). The sublingual ducts (Sd) opened directly into the sublingual mucosa (Figs. 3a-f); however, the mandibular ducts opened into the ventral and median side of the SC (Figs. 4a-f). The epithelium lining the submucosa was keratinized stratified squamous type, but it became non-keratinized at the opening of both ducts (Figs. 3 d-f, and 4d-f). Numerous fluorescently labelled Si beads were observed on the dorsal surface of the sublingual mucosa, especially that of the SC and near the opening of the sublingual ducts (Figs. 3a-f). Moreover, beads were observed in the lumina of both ducts (Figs. 3 e-f, and 4e-f). Interestingly, fluorescently labelled Si beads were observed in the apical cytoplasm of some cells lining the sublingual ducts (Figs. 3h-k), but not that in the mandibular ducts (Figs. 4e-f).

**Fig. 3.**
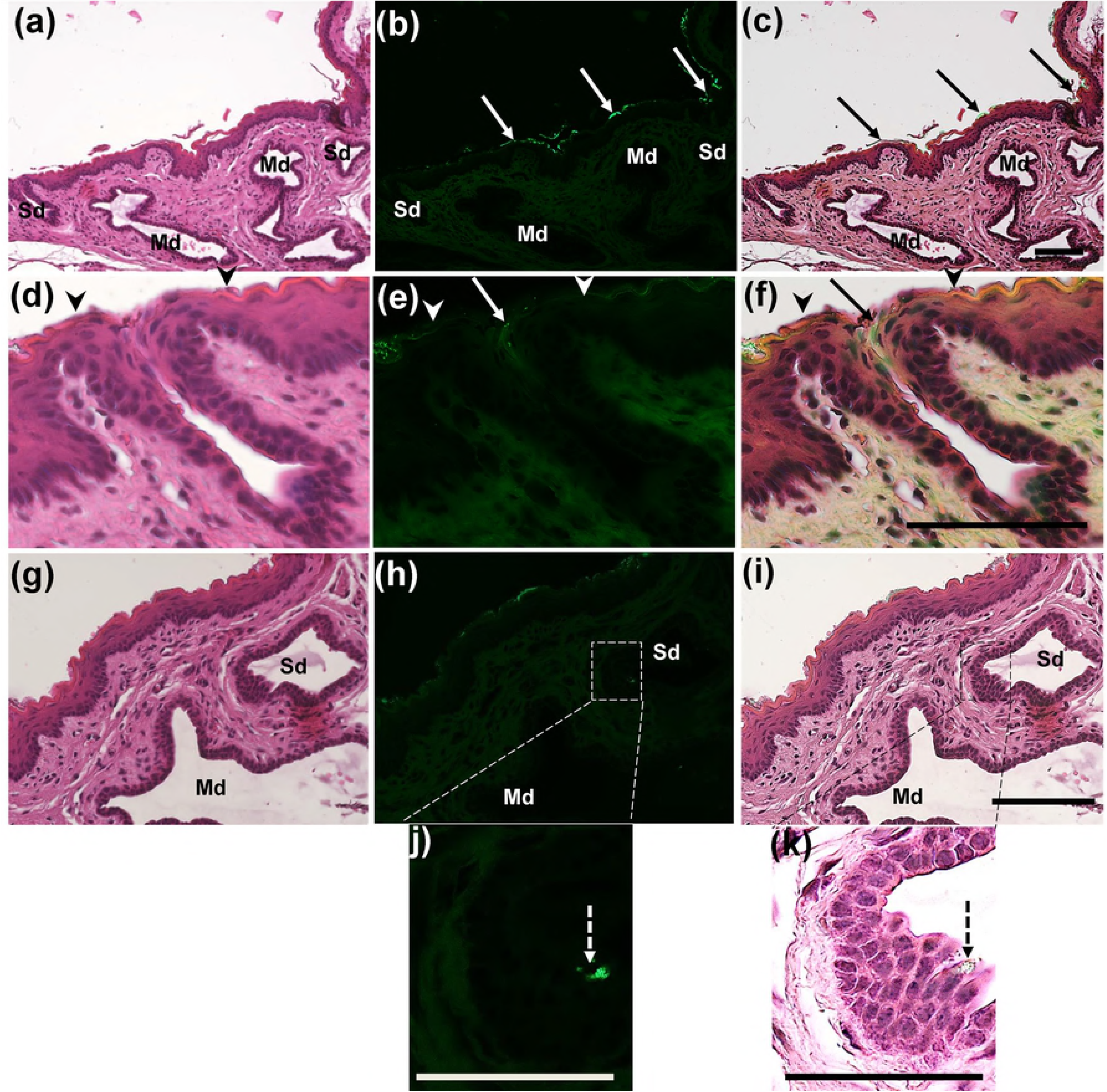
Fluorescence microscopic observation of the sublingual mucosa at the SC level. (a-c) Fluorescence microscopic images of H&E stained sections of the mandibular duct (Md) and the opening of the sublingual ducts (Sd) into the sublingual mucosa. Notice separate openings of the Md and Sd and numerous fluorescent beads accumulating (arrows) on the sublingual mucosa at the SC region and at the opening of the Sd. (d-f) Fluorescence microscopic images of H&E stained sections of the opening of the Sd. Notice the change of the keratinized epithelium (arrow heads) to non-keratinized type at the opening of the Sd (arrows). (g-i) The wall of the mandibular duct (Md) and the sublingual ducts (Sd). (j, k) Higher magnifications of the boxed areas in figure (h, i). Notice fluorescence beads in the apical cytoplasm of some ductal lining cells (dashed arrows).

**Fig. 4.**
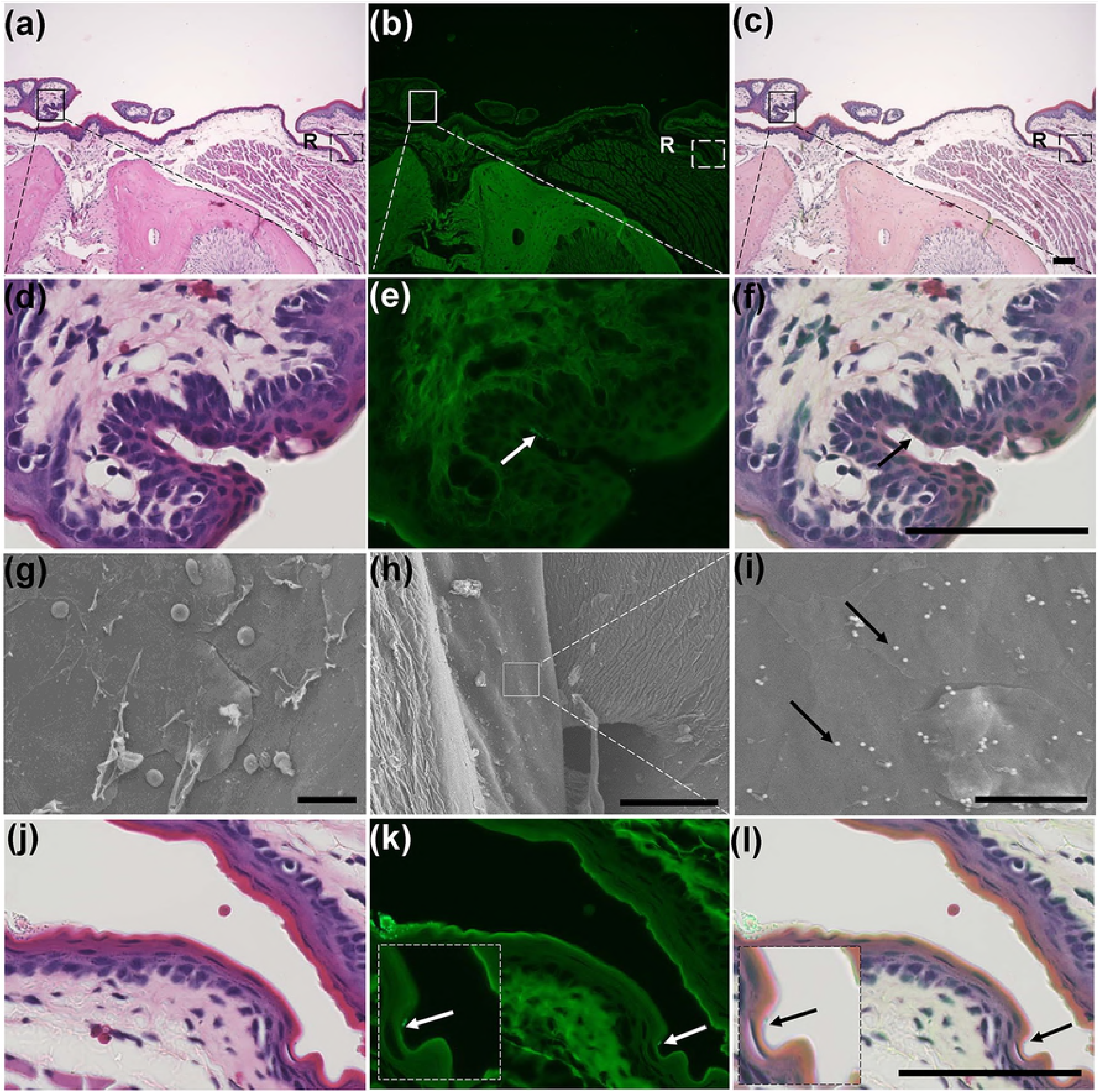
(a-c) Fluorescence microscopic observation of the sublingual mucosa at the levels of the BM and the opening of Md into the SC. Notice the opening of the Md into the SC (solid box) and the presence of a slight recess (R) representing the BM. (d-f) Higher magnifications of the boxed areas in figure (a-c). Notice the fluorescent beads in the lumen of the Md (arrows). (g-i) SEM of the dorsal surface of the BM of a 3-month-old female mouse of the control group (g), and the experimental group (h, i). Notice the smooth dorsal surface of the BM and several Si beads (arrows). (j-l) Higher magnifications of the dashed boxed areas in figure (a-c). Notice the fluorescent beads in the recess (arrows).

The SEM observation of the BM in both control (Fig. 4g) and Si bead group (Figs. 4h-i) revealed the smooth surface of the BM. In the experimental groups, Si beads could be observed on the dorsal surface of the BM, but fewer in number than that in the SC, OR (Fig. 4i). Examination of the H&E stained serial sections revealed a slight recess representing the BM (Figs. 4a-c) in which few Si beads were observed (Figs. 4j-l). The epithelium lining the BM was keratinized stratified but of thinner thickness the surrounding epithelium, and consisted of 2-3 layers (Figs. 4a-c, and 4j-l).

The SEM observation of the OR of the control group revealed a shallow median depression with a smooth dorsal surface and edge (Figs. 5a-c). In experimental groups, many Si beads were observed both on the dorsal surface and in the groove (Figs. 5d-f). In the H&E stained sections, the mucosa of the OR showed a shallow depression that was lined by keratinized stratified squamous epithelium of slightly thinner thickness than the surrounding mucosa. Few Si beads were observed attaching to the outer keratinized layer (Figs. 5g-l).

**Fig. 5.**
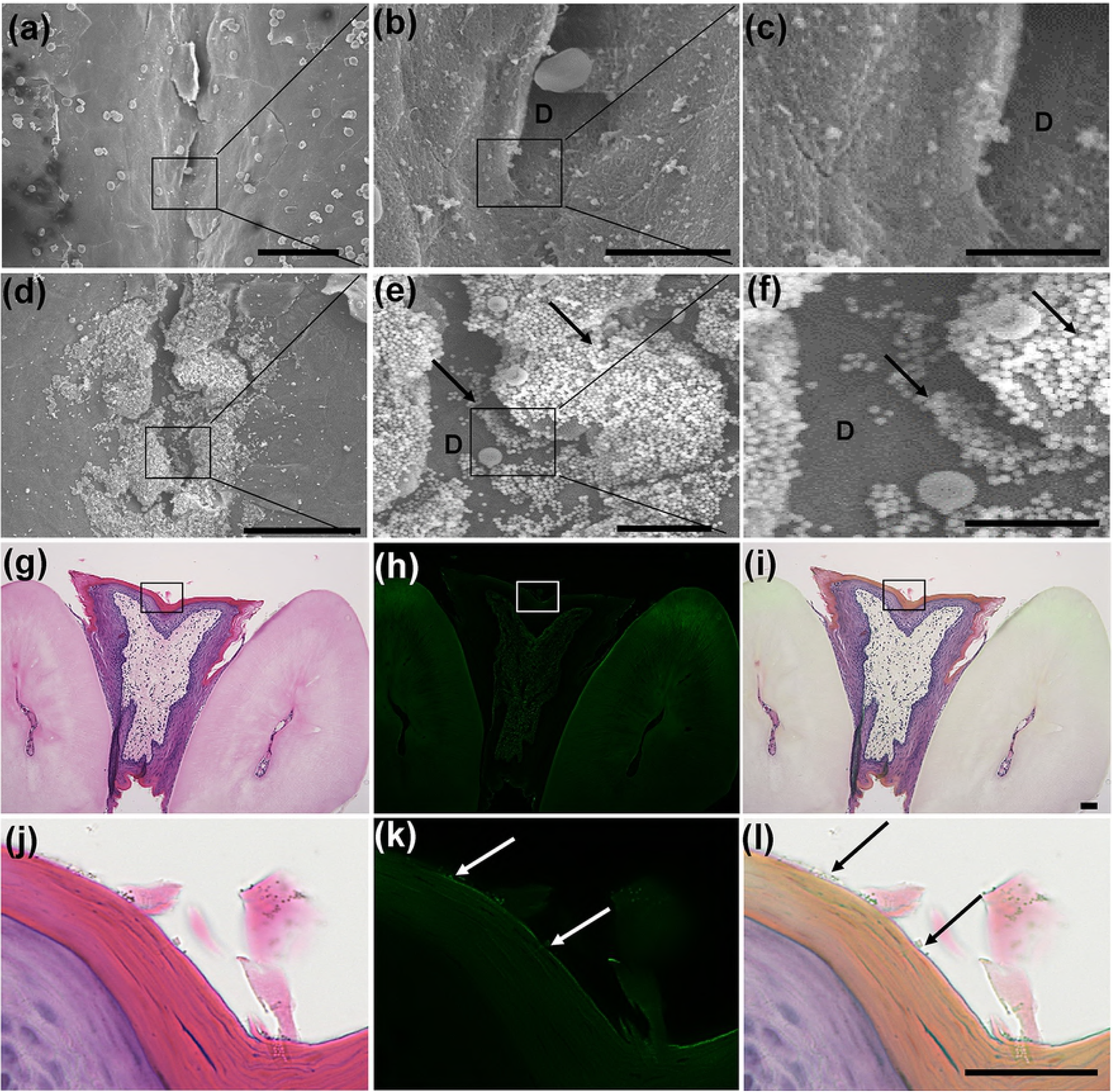
(a-f) SEM of the dorsal surface of the OR of a 3-month-old female mouse of the control group (a-c), and the experimental group (d-f). Notice a shallow depression (D) with a smooth dorsal surface and a smooth edge. A moderate number of beads were observed in the experimental group (arrows). (g-i) Fluorescence microscopic observation of the sublingual mucosa at the OR level. Notice the slight depression of the mucosa and decreased thickness of the lining epithelium compared to the surrounding epithelium. (j-l) Higher magnifications of the dashed boxed areas in figure (a-c). Notice the few fluorescent beads attached to the keratinized epithelium (arrows).

## DISCUSSION

SLIT is an effective and safe therapy that has recently been established as a valid method for the treatment of many allergic diseases via the induction of antigen-specific tolerance. It has been used to treat allergic diseases in humans such as allergic rhinitis, allergic rhinoconjunctivitis, and asthma [22-28], as well as those in animals, such as canine atopic dermatitis [2, 29]. Several approaches involving analysis of the immunological status of the oral mucosa have been used to elucidate the mechanism of antigen uptake following SLIT [13 -15]. After morphological analysis of the different compartments of the oral mucosa, their roles in antigen uptake remains unclear. Therefore, we undertook a detailed morphological characterization of the sublingual mucosa. This was achieved by administering fluorescent beads into the sublingual region to mimic antigen uptake and induce antigen-specific immune tolerance.

Our data revealed three main sites within the oral cavity where beads accumulated following their sublingual administration (SC, OR, and BM). These sites could reflect the locations where substances tend to stagnate anatomically or the sites where antigen can be up taken. Interestingly, our data revealed that within the same mice, beads accumulated in varying degrees in the three anatomic sites of the sublingual mucosa. The SC, OR, and BM showed higher, moderate, and lower tendency for accumulations, respectively. A previous report explained that there are three features of the oral epithelium (thickness, keratinization, and rete ridges) that could significantly alter allergen capture following sublingual allergen immunotherapy [30]. In support of this, our study revealed substantial bead accumulation on the dorsal surface of the SC. There was especially heavy near the central groove of the SC and the deep depression, representing the opening sites of the mandibular and sublingual ducts, respectively. Moreover, our SEM and light microscopic data revealed two features of the duct openings that could improve antigen uptake. First, there were abrupt changes in the surface of the duct openings from the surrounding smooth surface. Second, there was a transition of epithelium from keratinized to nonkeratinized at the site of duct openings.

The epithelium lining both the OR and BM were stratified squamous type but had a lower cell thickness than that of the surrounding. Additionally, in the OR site, a shallow depression was observed that could provide greater opportunity for antigen accumulation. In sum, our data suggested that the degree of bead accumulation at different sites could be due to variations in the local environment of the oral cavity, such as the keratinization and morphological variations.

In our investigation, we observed Si beads accumulating in the apical cytoplasm of some cells lining the sublingual ducts but not in the mandibular ducts. This observation is supported by a previous report concerning the role of the sublingual ductal system in incorporating and delivering sublingual antigens to ductal antigen-presenting cells [18]. Interestingly, in this previous report M cells were observed in the gastrointestinal mucosa and demonstrated the ability to take up antigen by phagocytosis [30, 31]. We believe that some cells within the epithelial lining of the sublingual duct phagocytosed beads into their apical cytoplasm. These could be considered “M-like cells,” and may play a major role in the antigen uptake mechanism. However, in previous studies some serum components such as albumin have been reported to migrate from capillaries to saliva via interstitial fluid [32, 33]. Despite the fact that the molecular weight of albumin (66 kDa) is greater than some allergens such as cedar pollen allergens (36 kDa), there have been no reports demonstrating that allergens migrate from saliva into blood. Therefore, some other mechanism besides antigen uptake may be responsible for the effectiveness of SLIT. Further investigations are required.

Interestingly, a recent comparative study revealed that rodents and especially mice, could be used as animal models for pharmacodynamics/efficacy studies of SLIT [17]. Therefore, another goal of our investigation was to examine whether age and sex affect the efficacy of SLIT in mice. We found no remarkable difference in mucosal surface morphology between sexes. Interestingly, our data showed some variations in Lt bead accumulation between sexes, but none in Si bead accumulation. This is despite the fact that both types of beads were of equal size. Notably, in the Lt bead group, females tended to exhibit more fluorescence intensity at all observation sites, and significant sex differences were observed in the SC. However, such sex differences were not observed in the Si bead group. Variations between these groups may be due to the surface structure of the beads and their interaction or adhesion with mucous membranes. Additionally, the contents of saliva differ between male and female mice. This is attributable to the sexual dimorphism of intervening ducts in the mandibular gland [20]. Thus, we believe that variations in the oral environments of male and female mice could affect adhesion or uptake of beads into mucous membranes. In particular, the local environmental of the female oral mucosa may contribute to the accumulation of latex beads.

In conclusion, our investigation revealed the major role of some sublingual ductal epithelial cells in the antigen uptake mechanism following SLIT. Furthermore, our data revealed possible sex-related differences in the efficacy of SLIT. Specifically, females demonstrated a greater tendency toward bead accumulation. However, further investigations are required. This may include examining the effect of cyclic changes in female mice on bead accumulation.

## Supporting information

S1 Fig. Morphological observation of sublingual mucosa in mouse lower jaw following PBS or beads administration. (a) Method of sublingual administration with a 20μl micropipette of either PBS or beads to the lower part of the mouse’s tongue after raising the tongue. (b) Fluorescent stereomicroscopic image of the sublingual mucosa after cutting the anterior end of the tongue in the experimental group (latex beads, 3-month-old males). Notice fluorescently labelled bead accumulations in the sublingual caruncle (SC), buccal mucosa (BM), and oral rostral (OR) in front of the incisor teeth (IT) with no fluorescence detected in the negative control area (CA). (c) SEM image of the previous areas. (d) Fluorescent stereomicroscopic image of the sublingual mucosa in the control group. Notice the absence of fluorescent staining. Scale bar = 1 mm.

## References

1. Sakurai Y, Nakamura, K. Teruya K, Shimada N, Umeda T, Tanaka H, et al. Prevalence and risk factors of allergic rhinitis and cedar pollinosis among Japanese men. Prev Med. 1998; 27: 617-622. https://doi.org/10.1006/pmed.1998.0336 PMID: 9672957

2. Nødtvedt A, Egenvall A, Bergvall K, Hedhammar A. Incidence of and risk factors for atopic dermatitis in a Swedish population of insured dogs. Vet Rec. 2006; 159: 241-246. http://dx.doi.org/10.1136/vr.159.8.241 PMID: 16921013

3. Diesel A. Cutaneous hypersensitivity dermatoses in the feline patient: a review of allergic skin disease in cats. Vet Sci. 2017; 4: 25. http://doi:10.3390/vetsci4020025 PMID: 29056684

4. Muñoz-Cano R, Pascal M, Araujo G, Goikoetxea MJ, Valero AL, Picado C, et al. Mechanisms cofactors and augmenting factors involved in anaphylaxis. Front Immunol. 2017; 8: 1193. https://doi:10.3389/fimmu.2017.01193 PMID: 29018449

5. Townley, RG, Suliaman F. The mechanism of corticosteroids in treating asthma. Ann Allergy. 1987; 58: 1-6. PMID: 3026210

6. Ciprandi G, Tosca M, Passalacqua G, Canonica GW, Ricca V, Landi M. Continuous antihistamine treatment controls allergic inflammation and reduces respiratory morbidity in children with mite allergy. Allergy. 1999; 54: 358-365. https://doi.org/10.1034/j.1398-9995.1999.00920.x PMID: 10371095

7. Dahl R, Kapp A, Colombo G, Monchy JGR, Rak S, Emminger W, et al. Sublingual grass allergen tablet immunotherapy provides sustained clinical benefit with progressive immunological changes over 2 years. J Allergy Clin Immunol. 2008; 121: 512-518. https://doi.org/10.1016/j.jaci.2007.10.039 PMID:18155284

8. Ott H, Sieber J, Brehler R, Fölster-Holst R, Kapp A, Klimek L, et al. Efficacy of grass pollen sublingual immunotherapy for three consecutive seasons and after cessation of treatment: the ECRIT study. Allergy. 2009; 64: 1394-1401. https://doi.org/10.1111/j.1398-9995.2008.01875.x PMID: 19764942

9. Zhong C, Yang W,Li Y, Zou L, Deng Z, Liu M, and Huang X. Clinical evaluation for sublingual immunotherapy with Dermatophagoides farinae drops in adult patients with allergic asthma. Ir. J. Med. Sci. 2017; 1-6. http://doi:10.1007/s11845-017-1685-x. PMID: 29032417

10. Jay DC, and Nadeau KC. Immune mechanisms of sublingual immunotherapy. Curr Allergy Asthma Rep. 2014; 14: 473. http://doi:10.1007/s11882-014-0473-1 PMID:25195100

11. Zielen S, Devillier P, Heinrich J, Richter H, Wahn U. Sublingual immunotherapy provides long-term relief in allergic rhinitis and reduces the risk of asthma: a retrospective, real-world database analysis. Allergy. 2018; 73: 165-177. http://doi:10.1111/all.13213 PMID: 28561266

12. DeBoer DJ, Verbrugge M, Morris M. Clinical and immunological responses of dust mite sensitive, atopic dogs to treatment with sublingual immunotherapy (SLIT). Vet Dermatol. 2016; 27: 82-e24. http://doi:10.1111/vde.12284 PMID: 26749020

13. Hovav A-H. Dendritic cells of the oral mucosa. Mucosal Immunol. 2014; 7: 27-37. http://doi:10.1038/mi.2013.42 PMID: 23757304

14. Mascarell L, Lombardi V, Louise A, Saint-Lu N, Chabre H, Moussu H, et al. Oral dendritic cells mediate antigen-specific tolerance by stimulating TH1 and regulatory CD4+ T cells. J Allergy Clin Immunol. 2008; 122: 603-609. https://doi.org/10.1016/j.jaci.2008.06.034 PMID: 18774396

15. Mascarell L, Rak S, Worm M, Melac M, Soulie S, Lescaille G, et al. Characterization of oral immune cells in birch pollen-allergic patients: impact of the oral allergy syndrome and sublingual allergen immunotherapy on antigen-presenting cells. Allergy. 2015; 70: 408-419. http://doi:10.1111/all.12576 PMID: 25631199

16. Zhang C, Ohno T, Kang S, Takai T, Azuma M. Repeated antigen painting and sublingual immunotherapy in mice convert sublingual dendritic cell subsets. Vaccine. 2014; 32: 5669-5676. http://doi:10.1016/j.vaccine.2014.08.013 PMID: 25168308

17. Thirion-Delalande C, Gervais F, Fisch C, Cuine J, Baron-Bodo V, Moingeon P, et al. Comparative analysis of the oral mucosae from rodents and non-rodents: application to the nonclinical evaluation of sublingual immunotherapy products. PLoS ONE. 2017; 12 (9): e0183398. https://doi.org/10.1371/journal.pone.0183398 PMID: 28886055

18. Nagai Y, Shiraishi D, Tanaka Y, Nagasawa Y, Ohwada S, Aso H, et al. Transportation of sublingual antigens across sublingual ductal epithelial cells to the ductal antigen-presenting cells in mice. Clin. Exper. Allergy. 2014; 45: 677-686. http://doi:10.1111/cea.12329 PMID: 24773115

19. Brook I. The bacteriology of salivary gland infections. Oral Maxillofac Surg Clin North Am. 2009; 21: 269-274. http://doi:10.1016/j.coms.2009.05.001 PMID: 19608044

20. Amano O, Mizobe K, Bando Y, Sakiyama K. Anatomy and histology of rodent and human major salivary glands. Acta Histochem Cytochem. 2012; 45: 241-250. http://doi:10.1267/ahc.12013 PMID: 23209333

21. Nagler RM. Salivary glands and the aging process: mechanistic aspects, health-status and medicinal-efficacy monitoring. Biogerontology. 2004; 5: 223-233. http://doi:10.1023/B:BGEN.0000038023.36727.50 PMID: 15314272

22. Wilson DR, Torres LI, Durham SR. Sublingual immunotherapy for allergic rhinitis. Cochrane Database Syst. Rev. 2003. CD002893. http://doi:10.1002/14651858 PMID: 12804442

23. Pajno GB, Peroni DG, Vita D, Pietrobelli A, Parmiani S, Boner AL. Safety of sublingual immunotherapy in children with asthma. Paediatr. Drugs. 2003; 5: 777-781. PMID: 14580226

24. Passalacqua G, Guerra L, Pasquali M, Lombardi C, Canonica GW.Efficacy and safety of sublingual immunotherapy. Ann. Allergy Asthma Immunol. 2004; 93: 3-12. http://doi:10.1016/S1081-1206(10)61440-8 PMID: 15281466

25. Penagos M, Compalati E, Tarantini F, Baena-Cagnani R, Huerta J, Passalacqua G, et al. Efficacy of sublingual immunotherapy in the treatment of allergic rhinitis in pediatric patients 3 to 18 years of age: a meta-analysis of randomized, placebo-controlled, double-blind trials. Ann. Allergy Asthma Immunol. 2006. 97: 141-148. http://doi:10.1016/S1081-1206(10)60004-X PMID: 16937742

26. Mascarell L, Van Overtvelt L, Moingeon P. Novel ways for immune intervention in immunotherapy: mucosal allergy vaccines. Immunol Allergy Clin North Am. 2006; 26: 283-306, vii-viii. http://doi:10.1016/j.iac.2006.02.009 PMID: 16701145

27. Dahl R, Kapp A, Colombo G, de Monchy JG, Rak S, Emminger W, et al. Efficacy and safety of sublingual immunotherapy with grass allergen tablets for seasonal allergic rhinoconjunctivitis. J Allergy Clin Immunol. 2006; 118: 434-440. http://doi:10.1016/j.jaci.2006.05.003 PMID: 16890769

28. Didier A, Malling HJ, Worm M, Horak F, Jager S, Montagut A, et al. Optimal dose, efficacy, and safety of once-daily sublingual immunotherapy with a 5-grass pollen tablet for seasonal allergic rhinitis. J Allergy Clin Immunol. 2007; 120: 1338-1345. http://doi:10.1016/j.jaci.2007.07.046 PMID: 17935764

29. Marsella R. Tolerability and clinical efficacy of oral immunotherapy with house dust mites in a model of canine atopic dermatitis: a pilot study. Vet. Dermatol. 2010; 21: 566-571. http://doi:10.1111/j.1365-3164.2010.00890.x. PMID: 20492623

30. Hase K, Ohno H. Epithelial cells as sentinels in mucosal immune barrier. Clin Immunol. 2006; 29: 16-26. http://doi:10.2177/jsci.29.16 PMID: 16505599

31. Kimura S. Molecular insights into the mechanisms of M-cell differentiation and transcytosis in the mucosa-associated lymphoid tissues. Anat Sci Int. 2017; 1-12. http://doi:10.1007/s12565-017-0418-6 PMID: 29098649

32. Vining FV, McGinley RA, Symons RG. Hormones in saliva: mode of entry and consequent implivations for clinical interpretation. Clin Chem. 1983; 29: 1752-1756. PMID: 6225566

33. Pfaffe T, Cooper-White J, Beyerlein P, Kostner K, Punyadeera C. Diagnostic potential of saliva: current state and future applications. Clin Chem. 2011; 57: 675-687. http://doi:10.1373/clinchem.2010.153767 PMID: 21383043

